# Circulation of third-generation cephalosporin resistant *Salmonella* Typhi in Mumbai, India

**DOI:** 10.1101/2021.08.25.457312

**Authors:** Silvia Argimón, Geetha Nagaraj, Varun Shamanna, Darmavaram Sravani, Ashwini Kodlipet Vasanth, Akshatha Prasanna, Aruna Poojary, Anurag Kumar Bari, Anthony Underwood, Mihir Kekre, Stephen Baker, David M. Aanensen, Ravikumar Kadahalli Lingegowda

**Affiliations:** Centre for Genomic Pathogen Surveillance, Wellcome Genome Campus, Hinxton, Cambridge, United Kingdom; Central Research Laboratory, Kempegowda Institute of Medical Sciences, Bengaluru, India; Department of Pathology and Microbiology, Breach Candy Hospital Trust, Mumbai, India; Cambridge Institute of Therapeutic Immunology & Infectious Disease, Department of Medicine, University of Cambridge, Cambridge, United Kingdom; Centre for Genomic Pathogen Surveillance, Li Ka Shing Centre for Health Information and Discovery, University of Oxford, Old Road Campus, Oxford, United Kingdom

**Author notes:** Corresponding author: Silvia Argimón, Centre for Genomic Pathogen Surveillance, Wellcome Genome Campus, Hinxton, United Kingdom. Alternate corresponding author: Geetha Nagaraj, Central Research Laboratory, Kempegowda Institute of Medical Sciences, Bengaluru, India. These authors contributed equally (Silvia Argimón, Geetha Nagaraj).

## Abstract

We report the persistent circulation of third-generation cephalosporin resistant *Salmonella* Typhi in Mumbai, linked to the acquisition and maintenance of a previously characterized IncX3 plasmid carrying the ESBL gene *bla*_SHV-12_ and the fluoroquinolone resistance gene *qnrB7* in the genetic context of a triple mutant also associated with fluoroquinolone resistance.

## Brief Report

Typhoid fever is a serious enteric disease spread through food and water contaminated with *Salmonella enterica* serovar Typhi (*S*. Typhi), which disproportionately affects populations in locations with limited sanitation and hygiene. There remains a significant burden of typhoid fever In India [1, 2], where antimicrobial therapy continues to be the linchpin of treatment and control. However, populations of *S*. Typhi rapidly develop resistance whenever a new antimicrobial is introduced for the treatment of typhoid fever [3].

Increasing rates of resistance against fluoroquinolones [4-7] linked to the expansion of genotype 4.3.2.1 (haplotype H58 lineage II) [8] have led to the adoption of cefixime and ceftriaxone (third generation cephalosporins) and azithromycin (macrolide) as the first-line treatments for typhoid fever in India [9]. This reliance on third generation cephalosporins may trigger the emergence of further resistance, as has been observed with extensively-drug resistant (XDR) *S*. Typhi in Pakistan [10, 11]. Third-generation cephalosporin resistant *S*. Typhi have been previously isolated in India and associated with the presence of the AmpC genes *bla*_CMY-2_ [12], *bla*_ACC-1_ [13], *bla*_DHA-1_ [14], and ESBL gene *bla*_SHV-12_ [15]. As with other Gram-negative bacteria, ESBL and AmpC genes in *S*. Typhi are often located on plasmids, which commonly also carry the fluoroquinolone resistance gene *qnr* [10, 15].

Despite the increasing use of third generation cephalosporins [3, 16], the prevalence of resistance in *S*. Typhi in India remains low, ranging from 0 to 5% [5, 7, 16-18]. However, a recent study from Mumbai reported a rate of ceftriaxone resistance of 12.5% [19].

To better understand the genetic context of third-generation cephalosporin resistant *S*. Typhi in Mumbai, we characterized the antimicrobial susceptibility profiles and the genomes of 92 isolates from blood culture-confirmed patients with *S*. Typhi infections collected by a tertiary care hospital in Mumbai between January 2017 and December 2018. The demographic and clinical characteristics of the patients are described in **Supplementary Table S1**. The species identification and antimicrobial susceptibility testing were performed using the VITEK-2 compact system and minimum inhibitory concentration (MIC) values were interpreted according to the Clinical Laboratory Standards Institute 2019 guidelines. Azithromycin and chloramphenicol were not included in the standard panel of antibiotics. All *S*. Typhi isolates were susceptible to piperacillin-tazobactam, ertapenem, imipenem, meropenem, and colistin. The majority (83/92; 90.2%) of the isolates were non-susceptible to ciprofloxacin. Twelve isolates (13.0%) were resistant to ceftriaxone, out of which 11/12 were also non-susceptible to cefepime. All isolates exhibiting phenotypic resistance to third-generation cephalosporins were also resistant to ciprofloxacin.

Two of the ceftriaxone-resistant isolates were recovered in 2017 while the remaining ten were recovered in 2018, mostly between January and July, suggesting a cluster of cases (**Supplementary Figure S1**). Out of 80 patients with clinical data, a higher proportion of patients with ceftriaxone-resistant *S*. Typhi were inpatients (8/11, 72.7%), compared to patients with ceftriaxone-susceptible *S*. Typhi (39/69, 56.2%), but this difference was not significant (Fisher exact test 0.51, *p* > 0.05).

To investigate the genetic relationship between the ceftriaxone-resistant isolates and to determine the molecular basis of resistance, we sequenced the 92 *S*. Typhi on Illumina HiSeq X10 with 150 bp paired-end reads, resulting in 89 whole genome sequences after assembly, quality control and analysis using pipelines developed within the National Institute for Health Research Global Health Research Unit on Genomic Surveillance of AMR [20]. Sequence data were deposited at the European Nucleotide Archive under study accession PRJEB29740. AMR genes and point mutations were identified from sequence reads using ARIBA v2.14.4 [21] with the NCBI database [22] and the pointFinder database [23]. Genotype [24] and plasmid replicon type [25] information, as well as AMR determinants, were obtained from assemblies using Pathogenwatch [26]. Details of the methods are provided in the supplementary materials.

Lineage 4.3.1 (haplotype H58) encompassed 87.6% of the genome sequences, with approximately three-quarters (67/89; 75.3%) of the genomes belonging to genotype 4.3.1.2, followed by 4.3.1.1 (6/89, 6.7%) and 4.3.1 (5/89, 5.6%). Resistance to ceftriaxone was associated with a *bla*_SHV-12_ ESBL gene in 11/12 isolates belonging to genotype 4.3.1.2. We did not identify any known mechanism of ceftriaxone resistance in the genome of the additional isolate, which was assigned to 4.3.1. This isolate was susceptible to ampicillin and the MIC for ceftriaxone was 4 mg/L (the breakpoint for ceftriaxone resistance), which were reproduced upon retesting. All isolates carried at least one mutation in the quinolone-resistance determining region in *gyrA*, but we did not identify any of the described mutations or genes that confer resistance to azithromycin. Only one organism was predicted to be MDR (i.e., resistant to ampicillin, chloramphenicol and sulfamethoxazole-trimethoprim), which belonged to genotype 4.3.1.1 and carried resistance genes typically located on an IncHI1 plasmid or in a composite transposon inserted into the chromosome [27] (**Figure 1A**). The lack of an IncHI1 replicon and further inspection of the genome assembly pointed to a *yidA* chromosomal insertion site.

**Figure 1.**
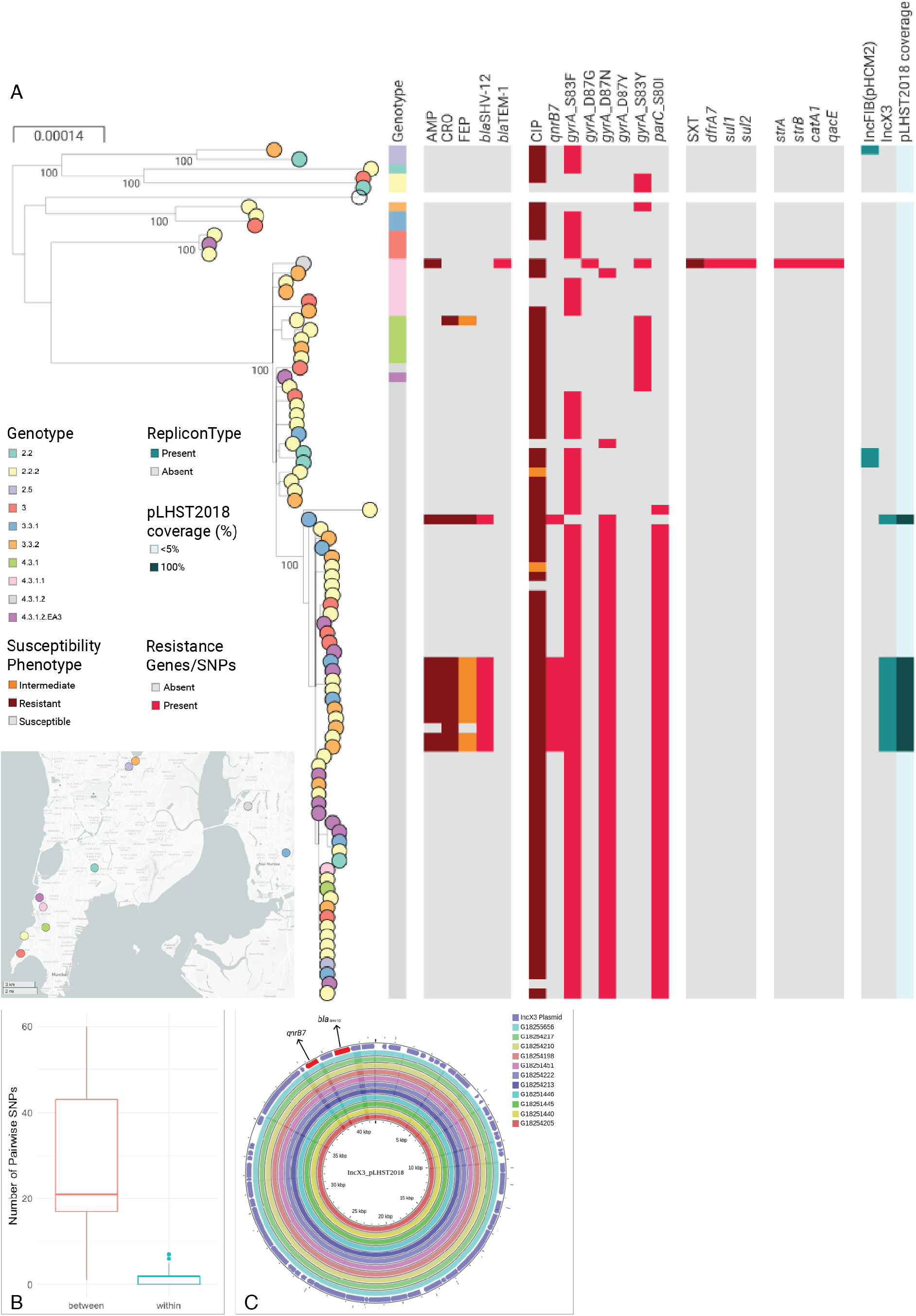
Analysis of the *S*. Typhi genomes. **A)** Phylogenetic tree of 89 genomes with nodes coloured by the neighbourhood in Mumbai where the patient resided. The maximum-likelihood tree was inferred from 1496 SNP positions identified by mapping the sequence reads to the complete chromosome of strain CT18 (NC_003198) and masking regions corresponding to mobile genetic elements and recombination from the alignment of pseudogenomes. The tree is annotated with the genotype; the susceptibility phenotype to ampicillin (AMP), ceftriaxone (CRO), cefepime (FEP), ciprofloxacin (CIP), and sulfamethoxazole-trimethoprim (SXT); the distribution of resistance genes and mutations conferring resistance to these antibiotics and others not tested; the distribution of plasmid replicon types identified in the genome assemblies with Pathogenwatch; the sequence coverage of the plasmid pLHST2018 determined by mapping sequence reads of each genome to the plasmid sequence. The data is available at https://microreact.org/project/S.Typhi_Mumbai_2017-2018/8713a72b. **B)** Boxplot showing the distribution of the pairwise SNP differences within the cluster of 10 isolates carrying *bla*_SHV-12_ (red) or between the genomes in this cluster and other 4.3.1.2 genomes not in this cluster (blue). The horizontal line indicates the median and the box indicates the interquartile range. **C)** Comparison of assembly contigs from 11 genomes in this study carrying *bla*_SHV-12_, *qnrB7* and the IncX3 replicon type to the complete sequence of plasmid pLHST2018. The outermost circle shows the plasmid genes, with resistance genes shown in red.

A phylogenetic investigation of the 89 genome sequences determined that the eleven 4.3.1.2 isolates harboring *bla*_SHV-12_ were located in two independent branches of the tree (**Figure 1A**). One group was represented by a single isolate characterized by the presence of *gyrA* mutation D87N, the *qnrB7* gene and the IncX3 replicon sequence. The second comprised ten isolates and also carried the *qnrB7* gene and the IncX3 type replicon, but was a triple mutant associated with fluoroquinolone resistance (D87N and S83F in *gyrA* and S80I in *parC*). This cluster of 10 isolates was supported by a 100% bootstrap value and a close genetic relationship (0-7 pairwise SNP differences) in comparison to the other 4.3.1.2 sequences (**Figure 1B**). An alignment of the assembly contigs from each of the eleven genomes carrying *bla*_SHV-12_, *qnrB7* and IncX3 to the complete sequence of the previously described plasmid pLHST2018 (accession CP052768 [15]) using the CGView Server [28] showed a complete match to the 42.8 Kbp plasmid sequence (**Figure 1C**). This was confirmed by mapping the sequence reads of each of the 89 genomes to the pLHST2018 plasmid and computing the sequence length coverage, which shows 100% coverage of the plasmid sequence only for the 11 genomes with *bla*_SHV-12_ (**Figure 1A**).

These data are consistent with two independent acquisitions of plasmid pLHST2018 within genotype 4.3.1.2, of which only one showed evidence of maintenance. The ten isolates in the cluster were recovered between April 2017 and December 2018 from patients residing in four different neighborhoods within a 30 km radius of the hospital. We compared the 67 4.3.1.2 genomes from this study to global public genomes from the same genotype using Pathogenwatch. Four previously described ceftriaxone-resistant isolates from Mumbai carrying *bla*_SHV-12_ in the IncX3 plasmid pLHST2018 [15] formed a monophyletic cluster and were interspersed in the tree with the isolates from this study (**Supplementary Figure S2**), signifying that this lineage has been circulating since at least 2016. Our data suggests that the persistence of the IncX3 plasmid conferring ceftriaxone resistance is not as short-term as previously thought [15], coinciding with the acquisition of the ceftriaxone resistance within the genetic context of the fluoroquinolone-resistant triple mutant lineage [8, 29].

The extended temporal and geographical distribution of the isolates suggests broad environmental transmission. Transmission from contaminated food or water at a public gathering, with subsequent person-to-person transmission within neighborhoods is also plausible. The higher proportion of male patients observed amongst ceftriaxone-resistant isolates (9/12, 75.0%) compared to ceftriaxone-susceptible isolates (34/80, 42.5%) is notable in this context, but not significant (Fisher exact test 0.06, *p* > 0.05, **Supplementary Table S1**). The absence of an epidemiological investigation based on patient exposures limits our understanding of the circulation of this high-risk clone. Although the isolates analyzed were collected by only one hospital in Mumbai, patients from all over the city attend the hospital and our lineage 4.3.1.2 S. Typhi genomes were interspersed with those from previous studies in the tree, suggesting adequate representation.

The World Health Organization classified third, fourth and fifth generation cephalosporins as critically important for human [30]. We characterized an emergent high-risk lineage of *S*. Typhi with plasmid-borne ceftriaxone and ciprofloxacin resistance in Mumbai, nested successively within a high-risk clone of chromosomally-encoded ciprofloxacin resistance found in several countries in South Asia and associated with fluoroquinolone treatment failure [29], the 4.3.1.2 genotype that has been expanding clonally in India [8], and the epidemic drug-resistant clone 4.3.1 clade or haplotype H58 [27]. The evidence of persistent transmission of this high-risk lineage calls for active surveillance and effective prevention measures to avert a large outbreak [31] and global dissemination [32, 33]. Through technology transfer and capacity building, the Central Research Laboratory at KIMS, Bangalore, is now equipped to carry out whole genome sequencing [34] and bioinformatics analysis [35] locally to support genomic surveillance.

## Acknowledgements

We are grateful to the DNA Pipelines and Pathogen Informatics teams at the Wellcome Sanger Institute for their support.

