## Supplementary Materials for "Circulation of third-generation cephalosporin resistant *Salmonella* Typhi in Mumbai, India"

**Supplementary Table S1. Epidemiological and clinical characteristics of 92 patients with *S.* Typhi infection, discriminated by their susceptibility to ceftriaxone.**

| Variable | Total Isolates (*n* = 92) | Susceptible to Ceftriaxone (*n* = 80) | Resistant to Ceftriaxone (*n* = 12) |
|---|---|---|---|
| Patient age (in years), median (range) | 29 (3-68) | 29.5 (10-68) | 27.5 (3-56) |
| Patient sex, *n*(%) | | | |
| Female | 49 (53.2) | 46 (57.5) | 3 (25.0) |
| Male | 43 (46.7) | 34 (42.5) | 9 (75.0) |
| Hospitalization, *n* (%[a]) | | | |
| Inpatients | 47 (58.8) | 39 (56.5) | 8 (72.7) |
| Outpatients | 33 (41.2) | 30 (43.5) | 3 (27.3) |
| No data | 12 | 11 | 1 |
| Ceftriaxone MICs (in mg/L), range | <=1-32 | <=1 | 4-32 |

[a] Percent of the number of patients with hospitalization data for each column.

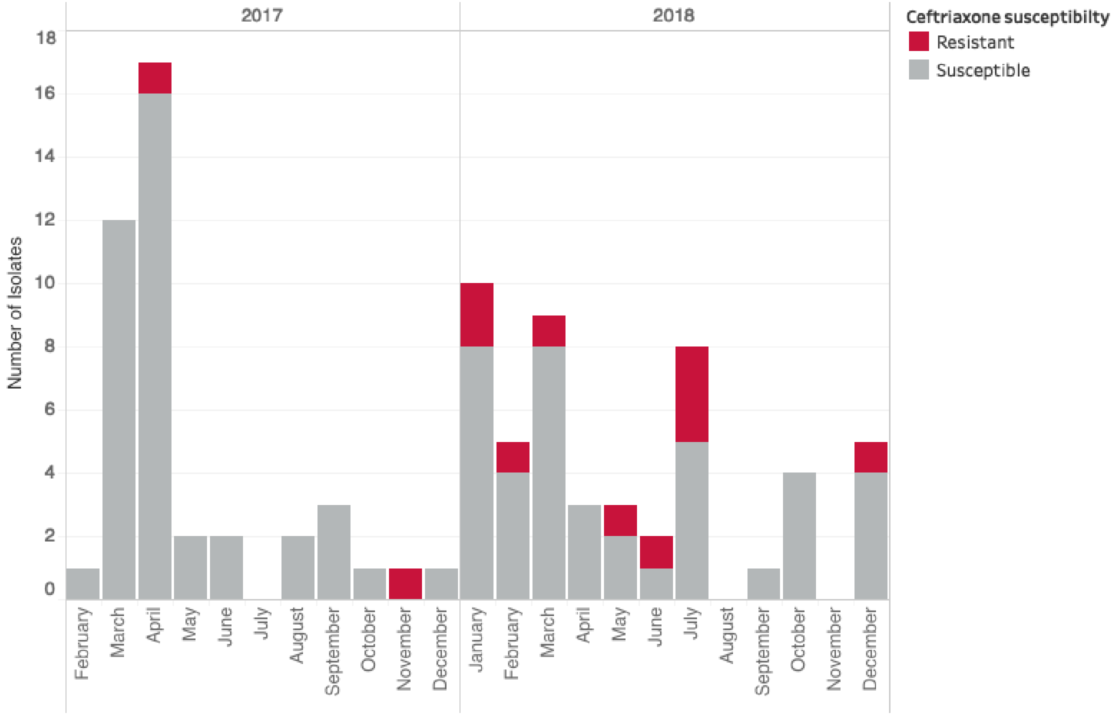


**Supplementary Figure S1. Temporal distribution of 92 *S.* Typhi isolates from Mumbai**.

The stacked bars were coloured by the isolates' susceptibility to ceftriaxone.

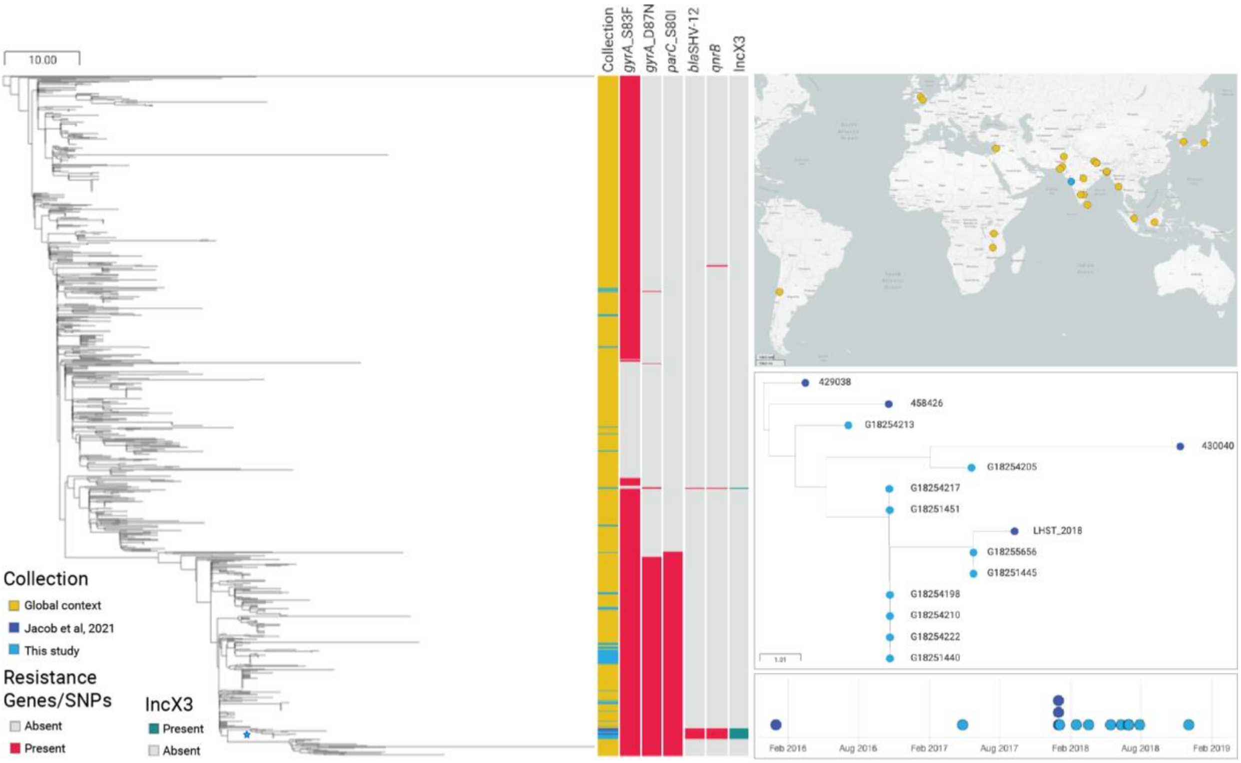


**Supplementary Figure S2. Pathogenwatch tree of genomes belonging to genotype 4.3.1.2.** Genomes from this study and from Jacob et al, 2021 [1] were contextualized with 816 additional genomes belonging to 4.3.1.2 using Pathogenwatch. The map shows the global distribution of the genomes coloured by collection. The node indicated with a star on the tree and comprising genomes with *bla*$_{SHV-12}$ is shown in detail in the tree panel on the right, together with the isolation timeline. The Pathogenwatch collection can be explored at [https://pathogen.watch/collection/j9873uuh3zw2-4312](https://pathogen.watch/collection/j9873uuh3zw2-4312)

### Supplementary Methods

*Bacterial strains*

A total of 92 *S.* Typhi isolates were obtained from a 213-bed tertiary care hospital in Mumbai, a densely populated city situated on the west coast of India with a population of 12.5 million inhabitants. The hospital is situated on the coastline of South Mumbai and caters to approximately 57,000 outpatients per year, as well as acting as a reference for antimicrobial susceptibility testing for the Mumbai area. The isolates were collected from patients with bloodstream infections suffering from enteric fever during 2017-2018 as a part of routine diagnosis and management of infectious disease activity of the hospital. Associated patient demographic and epidemiological data were also obtained. The study was approved by the KIMS Institutional Ethics Committee (KIMS/IEC/S12-2017, dated February 15th, 2018), and the Breach Candy Hospital Ethics Committee (BCMRC/P3/2018 dated February 7$^{th}$, 2018).

*Antimicrobial susceptibility testing*

The species identification and antimicrobial susceptibility testing were performed using the VITEK-2 compact system (Biomeurieux). Minimum inhibitory concentration (MIC) values were interpreted according to the Clinical Laboratory Standards Institute (CLSI) 2019 guidelines [2].

*Whole-genome sequencing and analysis*

Genomic DNA was isolated from 92 purified bacterial isolates using the Qiagen QIAamp DNA Mini Kit according to the manufacturer’s instructions. Double-stranded DNA libraries with 450 bp insert size were prepared and sequenced on the Illumina HiSeq X10 platform with 150 bp paired-end chemistry.

The sequence data were assembled using the pipeline developed within the National Institute for Health Research Global Health Research Unit on Genomic Surveillance of AMR (GHRU-AMR) [3] with the SPAdes assembler v3.14 [4], and subsequently annotated with

Prokka v1.5 [5]. Quality control parameters output by this pipeline included total assembly size, N50 score (> 25000), contaminant level (< 5%), and number of contigs (< 300). Three genomes were excluded based on the high number of heterozygous single-nucleotide polymorphisms (SNPs) detected when mapping sequence reads to the reference genome sequence of *S.* Typhi strain CT18 (NC_003198).

AMR genes and point mutations were identified in the genomes from sequence reads using ARIBA v2.14.4 [6] with the NCBI database and the pointFinder database adapted for ARIBA [3]. The genome assemblies were analyzed with Pathogenwatch [7], which predicts genotype (MLST, GenoTyphi genotype), plasmid replicon type, AMR profile, and presence of AMR determinants.

To characterize the insertion site of a putative composite transposon in one genome predicted to be MDR, the assembly contigs were ordered using abacas 1.3.1[8] against the reference genome ERL12148 (accession LT883153.1) [9] and compared using the Artemis Comparison Tool (ACT) [10] to visualise the genomic context of contigs corresponding to the MDR composite transposon.

A single-nucleotide polymorphism SNP-based phylogeny was generated with the pipeline developed by GHRU-AMR [11] by mapping reads to the complete chromosome sequence of *S.* Typhi strain CT18 (accession NC_003198) using the Burrow-Wheeler Aligner bwa mem v0.7.17[12]. Variants were called and filtered using bcftools v1.9 [13]. The mobile genetic elements and repetitive sequences in the genome of CT18 previously defined [14, 15] were masked in the pseudo-genome alignment using MGEmasker [16]. Recombination regions were removed using Gubbins v2.0.0 [17] and the non-recombinant SNPs were used to infer a maximum-likelihood tree using IQ-tree v1.6.8 [18] with the generalized time reversible model with a gamma correction for among site rate variation (GTR+G) and 100 bootstrap replicates, and with parameters -czb to collapse near zero branches. The tree was midpoint rooted and visualized in Microreact [19] along with the isolate metadata and the results of genomic analyses (https://microreact.org/project/S.Typhi_Mumbai_2017-2018). The alignment of non-recombinant SNPs was also used to compute the pairwise SNP differences between genomes with FastaDist v1.0.1 [20].

Genomes were analyzed further to determine the presence of plasmid pLHST2018 (accession CP052768) [1]. First, the assembly contigs of each of the 11 genomes harbouring the *bla*$_{SHV-12}$ gene and the IncX3 sequence were compared to the pLHST2018 plasmid sequence via a blastn comparison with default parameters on the CGView Server v1.0 [21]. Second, the sequence reads of each of the 89 genomes were mapped to the complete sequence of plasmid pLHST2018 with bwa mem and sequence coverage was computed with bedtools v2.29.2 [22] using the coverage assessment pipeline developed by GHRU-AMR [23].

We contextualized the 67 4.3.1.2 genomes in this study with four genomes from a recent study [1] and 816 public genomes also from this genotype available in Pathogenwatch. A neighbour joining tree was inferred from a matrix of pairwise SNP differences detected as described in detail previously [7] (Argimón et al. 2021). The Pathogenwatch collection is available at https://pathogen.watch/collection/j9873uuh3zw2-4312. A Microreact visualization was then created, which is available at https://microreact.org/project/Global_4-3-1-2.

https://www.protocols.io/view/ghru-genomic-surveillance-of-antimicrobial-resista-bpn6mmhe.

4. Bankevich A, Nurk S, Antipov D, et al. SPAdes: a new genome assembly algorithm and its applications to single-cell sequencing. J Comput Biol **2012**; 19(5): 455-77.
5. Seemann T. Prokka: rapid prokaryotic genome annotation. Bioinformatics **2014**; 30(14): 2068-9.
6. Hunt M, Mather AE, Sanchez-Buso L, et al. ARIBA: rapid antimicrobial resistance genotyping directly from sequencing reads. Microb Genom **2017**; 3(10): e000131.
7. Argimon S, Yeats CA, Goater RJ, et al. A global resource for genomic predictions of antimicrobial resistance and surveillance of *Salmonella* Typhi at pathogenwatch. Nat Commun **2021**; 12(1): 2879.
8. Assefa S, Keane TM, Otto TD, Newbold C, Berriman M. ABACAS: algorithm-based automatic contiguation of assembled sequences. Bioinformatics **2009**; 25(15): 1968-9.
9. Wong VK, Baker S, Pickard DJ, et al. Phylogeographical analysis of the dominant multidrug-resistant H58 clade of *Salmonella* Typhi identifies inter- and intracontinental transmission events. Nat Genet **2015**; 47(6): 632-9.
10. Carver T, Berriman M, Tivey A, et al. Artemis and ACT: viewing, annotating and comparing sequences stored in a relational database. Bioinformatics **2008**; 24(23): 2672-6.
11. Underwood A. SNP phylogeny nextflow pipeline. Available at: https://gitlab.com/cgps/ghru/pipelines/snp_phylogeny.
12. Li H, Durbin R. Fast and accurate long-read alignment with Burrows-Wheeler transform. Bioinformatics **2010**; 26(5): 589-95.
13. samtools. bcftools. Available at: https://github.com/samtools/bcftools.
14. Holt KE, Parkhill J, Mazzoni CJ, et al. High-throughput sequencing provides insights into genome variation and evolution in *Salmonella* Typhi. Nat Genet **2008**; 40(8): 987-93.

15. Ingle DJ, Nair S, Hartman H, et al. Informal genomic surveillance of regional distribution of *Salmonella* Typhi genotypes and antimicrobial resistance via returning travellers. PLoS Negl Trop Dis **2019**; 13(9): e0007620.

16. Underwood A. MGEmasker. Available at: https://gitlab.com/antunderwood/mgemasker.

17. Croucher NJ, Page AJ, Connor TR, et al. Rapid phylogenetic analysis of large samples of recombinant bacterial whole genome sequences using Gubbins. Nucleic Acids Res **2015**; 43(3): e15.

18. Nguyen LT, Schmidt HA, von Haeseler A, Minh BQ. IQ-TREE: a fast and effective stochastic algorithm for estimating maximum-likelihood phylogenies. Mol Biol Evol **2015**; 32(1): 268-74.

19. Argimón S, Abudahab K, Goater RJ, et al. Microreact: visualizing and sharing data for genomic epidemiology and phylogeography. Microb Genom **2016**; 2(11): e000093.

20. Underwood A. FastaDist. Available at: https://gitlab.com/antunderwood/fastadist.

21. Grant JR, Stothard P. The CGView Server: a comparative genomics tool for circular genomes. Nucleic Acids Res **2008**; 36(Web Server issue): W181-4.

22. Quinlan AR, Hall IM. BEDTools: a flexible suite of utilities for comparing genomic features. Bioinformatics **2010**; 26(6): 841-2.

23. Underwood A. Coverage assessment nextflow pipeline. Available at: https://gitlab.com/cgps/ghru/pipelines/dsl2/pipelines/coverage_assessment/-/tree/master/.
